# Unsupervised manifold alignment for single-cell multi-omics data

**DOI:** 10.1101/2020.06.13.149195

**Authors:** Ritambhara Singh, Pinar Demetci, Giancarlo Bonora, Vijay Ramani, Choli Lee, He Fang, Zhijun Duan, Xinxian Deng, Jay Shendure, Christine Disteche, William Stafford Noble

## Abstract

Integrating single-cell measurements that capture different properties of the genome is vital to extending our understanding of genome biology. This task is challenging due to the lack of a shared axis across datasets obtained from different types of single-cell experiments. For most such datasets, we lack corresponding information among the cells (samples) and the measurements (features). In this scenario, unsupervised algorithms that are capable of aligning single-cell experiments are critical to learning an *in silico* co-assay that can help draw correspondences among the cells. Maximum mean discrepancy-based manifold alignment (MMD-MA) is such an unsupervised algorithm. Without requiring correspondence information, it can align single-cell datasets from different modalities in a common shared latent space, showing promising results on simulations and a small-scale single-cell experiment with 61 cells. However, it is essential to explore the applicability of this method to larger single-cell experiments with thousands of cells so that it can be of practical interest to the community. In this paper, we apply MMD-MA to two recent datasets that measure transcriptome and chromatin accessibility in ~2000 single cells. To scale the runtime of MMD-MA to a more substantial number of cells, we extend the original implementation to run on GPUs. We also introduce a method to automatically select one of the user-defined parameters, thus reducing the hyperparameter search space. We demonstrate that the proposed extensions allow MMD-MA to accurately align state-of-the-art single-cell experiments.

## 1 Introduction

Most single-cell genomics assays measure a single property of the genome: scRNA-seq for mRNA expression, scMethyl-seq for methylation, scATAC-seq for chromatin accessibility, etc. The result is a detailed, single-cell view of the genome, but one in which only one property of each single cell can be measured. In some cases, pairs or even triplets of experimental assays can be combined into a single co-assay to explicitly measure multiple properties of the same cell (reviewed in [1]), but in general these co-assays tend to be lower throughput, more expensive, and noisier than the individual assays. Accordingly, a problem of increasing importance is to develop computational methods capable of integrating multiple types of single-cell measurements.

This general integration problem can be approached in two distinct ways: embedding or matching (Figure 1). In either case, a single cellular population is divided into two or more aliquots, and each aliquot is subjected to a different single-cell assay. The result is a collection of matrices in which rows correspond to individual cells and columns correspond to features of that cell. For scRNA-seq, the features are genes; for scATAC-seq the features are peaks, and for scMethyl-seq, the features are CpGs. Notably, all of the axes are disjoint: no cells and no features are shared across the different data sets. In the matching approach, for each pair of data sets, cells are matched to another in a one-to-one fashion with the aim of matching cells that share similar physical properties. In the embedding approach, integration is accomplished by projecting all of the cells into a shared latent space, usually of some user-specified dimensionality *p*. Cells that are close to one another in this latent space are inferred to share similar properties. In the machine learning literature, the problem of finding such a shared latent space is referred to as *manifold alignment* [2]. Note that any embedding method can be adopted to also solve the matching task, simply by carrying out a matching procedure in the latent space. Conversely, it is possible to use a matching to induce an embedding [3].

**Figure 1:**
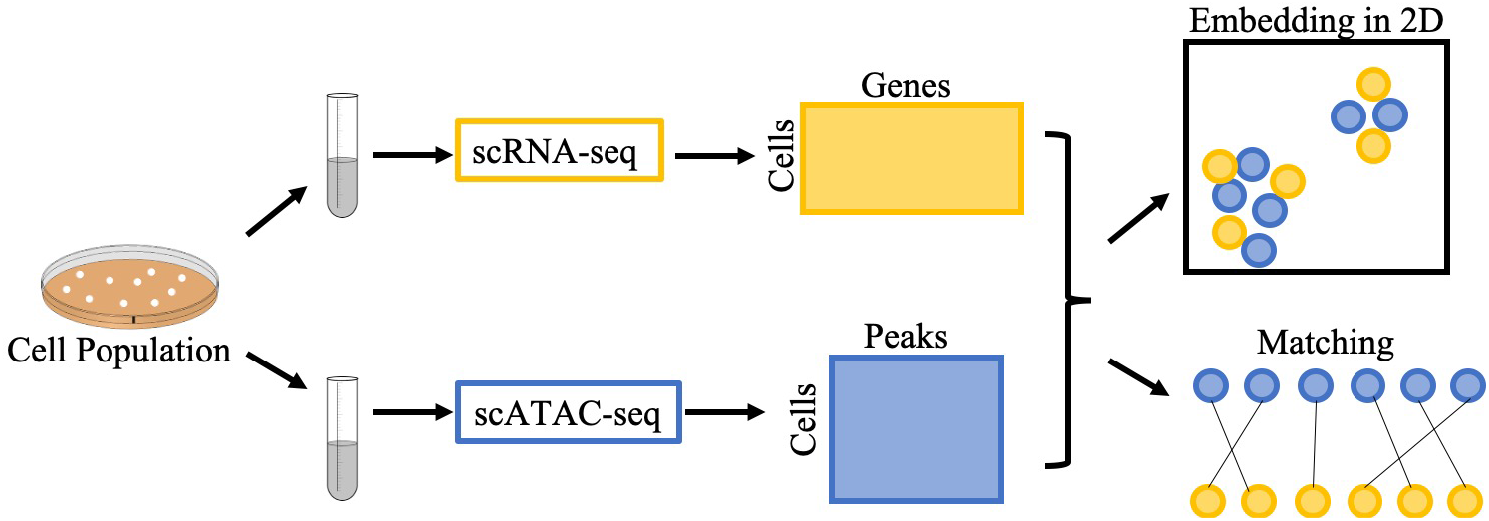
Two approaches to integration of heterogeneous single-cell data. A population of cells is divided into aliquots and subjected to two different single-cell assays. The measurements for each cell can either be embedded into a shared latent space (top right) or explicitly matched between data sets (bottom right).

A variety of algorithms have been developed to address both variants of this integration problem (Table 1). For embedding, some algorithms in the machine learning literature predate the existence of singlecell genomics data. For example, the joint Laplacian manifold alignment (JLMA) algorithm [2] constructs a cross-domain Laplacian submatrix using *k*-NN graphs that capture the local geometry of the data and then performs eigenvalue decomposition to find an embedding. One of the earliest manifold alignment methods explicitly developed for single-cell data, MATCHER [5], uses a Gaussian process latent variable model to project two data sets onto a 1D “pseudotime” trajectory. The model outputs a matching computed in this 1D space. More recently, LIGER [7] integrates single-cell data using an integrative non-negative matrix factorization to project the data into a shared low-dimensional space and then performs joint clustering in this space using a neighborhood graph. Finally, the latest version of the Seurat software (v3) [8] aligns pairs of single-cell data from different modalities but matching feature space by using canonical correlation analysis with *l*_2_-normalization.

**Table 1:** Algorithms for aligning multi-omic single-cell data.

| Name | Year | Approach | Problem | Supervised |
| --- | --- | --- | --- | --- |
| JLMA [2] | 2011 | Eigenvalue decomposition on joint Laplacian | Embedding | No |
| GUMA [4] | 2014 | Optimize geometry matching and feature matching | Matching | No |
| MATCHER [5] | 2015 | Gaussian process to 1D trajectory | Embedding | No |
| MAGAN [6] | 2018 | Generative adversarial network | Matching | Yes |
| LIGER [7] | 2019 | Non-negative matrix factorization | Embedding | Yes |
| Seurat v3 [8] | 2019 | Canonical correlation analysis | Embedding | Yes |
| MMD-MA [9] | 2019 | Optimize maximum mean discrepancy | Embedding | No |
| UnionCom [3] | 2020 | Optimize geometry matching and global scaling | Embedding | No |

The earliest method we are aware of for constructing an explicit matching between data sets is the generalized unsupervised manifold alignment (GUMA) algorithm [4], which optimizes an objective function with three terms: a geometry matching term across two different domains, a feature matching term, and a geometry preserving term for each domain. The algorithm aims to learn two matchings: not just a one-to-one correspondence between cells but also a one-to-one correspondence between features in the two domains. UnionCom [3] finds a matching between data sets using just the first matching term from GUMA. The optimization is extended to better handle datasets of different sizes, and non-linearity is introduced by running the matching on geodesic distances on a *k*-NN graph within each data set. Subsequently, the matching can be used to induce an embedding via a modified t-distribution Stochastic Neighbor Embedding (t-SNE) method. Another manifold alignment method, MAGAN [6], uses generative adversarial networks (GANs) to create a mapping between two different domains. Each domain is input into the generator of a GAN, which tries to produce the other domain as output. The generator aims to output samples such that the GAN’s discriminator module is unable to distinguish them from the real samples of the other domain. Thus, rather than creating a shared manifold, MAGAN produces two manifolds with an explicit alignment between them.

A key feature of each of these algorithms is whether the integration requires supervision in the form of *correspondence information*, i.e., information about which cells (rows) or features (columns) are matched. Of course, complete and perfect knowledge of the cell correspondence information is simply the solution to the mapping problem, so the correspondence information is typically incomplete (i.e., it is known for only a subset of cells or features) or noisy. JLMA, GUMA, MATCHER, and UnionCom are fully unsupervised algorithms. MAGAN can be run in unsupervised mode, but empirical results indicate that including some feature correspondence information is critical for good performance [6]. LIGER requires some feature-level correspondence information, since this is necessary to construct its neighborhood graph. Seurat also requires feature-level correspondence to perform the CCA step.

The current work builds upon a recently described method—maximum mean discrepancy manifold alighment (MMD-MA)—that approaches integration of heterogeneous single-cell data sets as an unsupervised embedding problem [9]. MMD-MA employs an objective function that minimizes the maximum mean discrepancy (MMD) between the data sets in the latent space, while also maintaining the underlying structure of each data set. The original paper describing MMD-MA demonstrates the excellent performance of the algorithm on three simulated data sets, as well as one real data set consisting of gene expression and methylation profiles of 61 single cells. Here, we provide an open source, GPU-based implementation of MMD-MA that allows it to scale up to the analysis of thousands of cells. We also demonstrate the utility of MMD-MA on two additional data sets. A co-assay data set, comprised of measurements of mRNA expression and chromatin accessibility, provides a gold standard by which we can explictly measure the quality of the matching induced by MMD-MA. We also apply MMD-MA to a differentiation time course dataset comprised of scRNA-seq and scATAC-seq measurements over five developmental time points of mouse embryonic stem cells (mESCs), demonstrating the algorithm’s ability to correctly match cells to neighbors from the same time point. As part of our optimization, we demonstrate how to reparameterize MMD-MA, effectively eliminating one of its key hyperparameters. And we provide empirical evidence that the GPU-based implementation provides a factor of 25 speedup relative to the original CPU implementation.

## 2 Methods

### 2.1 The maximum mean discrepancy manifold alignment algorithm

MMD-MA [9] optimizes an objective that embeds two datasets into a shared latent space. Let the two sets of points, 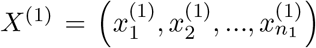, from 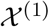 and 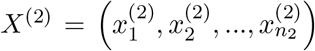, from 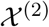, represent the two different single-cell measurements.

The input to the algorithm is in the form of similarity matrices *K_1_* and *K_2_* generated for domain 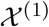 and 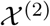, respectively. The pairwise similarities between the samples in a domain are calculated using positive definite kernel functions 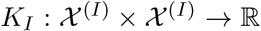 for *I* =1,2. This formulation makes the method generalizable across any type of data, like vectors, graphs, or strings, for which a kernel function can be defined. In this work, we use a scalar product kernel for both *K*_1_ and *K*_2_.

Mapping matrices *α*_1_ and *α*_2_, of size *n*_1_ × *p* and *n*_2_ × *p* respectively, project the samples from the two domains into a shared latent space of dimension *p*. MMD-MA optimization learns these mappings such that the projections *K*_1_*α*_1_ and *K*_2_*α*_1_ are as similar as possible. Specifically, MMD-MA optimizes, with respect to *α*_1_ and *α*_2_, an objective function that consists of three components. The first component is the MMD term to encourage the differently measured points to have similar distributions in the latent space:

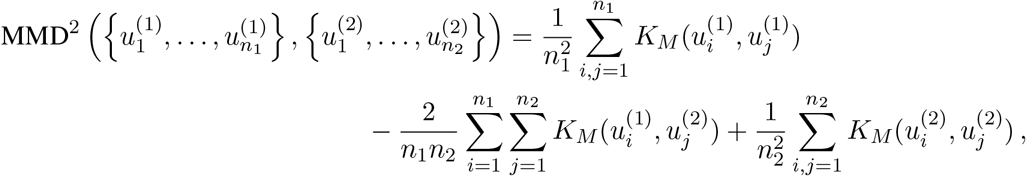

where 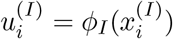, and *ϕ_I_* is the mapping of the input space to the *p*-dimensional latent space. Following Liu *et al*. [9], we use a radial basis function (RBF) kernel for *K_M_*:

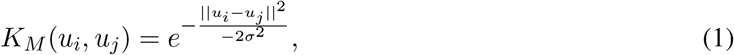

where *σ* is the bandwidth parameter. The second component of the MMD-MA objective is a distortion term to preserve the structure of the data between the input space and the latent space:

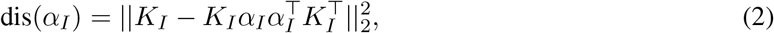

where *I_p_* is an identity matrix such that *p* is the number of learned features for the dataset. The third component avoids collapse to a trivial solution:

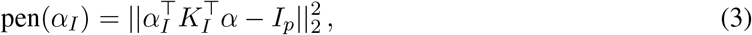

Thus, the final objective is

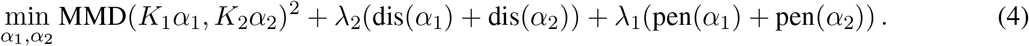

The MMD-MA algorithm requires that the user specify four hyperparameters: the dimensionality *p* of the latent space, the relative weights λ_1_ and λ_2_ of the second and third terms in the objective (Equation 4), and the bandwidth parameter *σ* for the RBF kernel in the MMD term.

### 2.2 Reparameterization of the kernel bandwidth

As with any kernel function, the RBF kernel *K_M_* in Equation 1 implicitly projects the data into a reproducing kernel Hilbert space [10]. Although, in principle, the Hilbert space induced by an RBF kernel is infinite dimensional, in practice any finite data set resides in a finite subspace of the full space. The bandwidth parameter *σ* associated with the kernel corresponds to the width of a Gaussian that is placed over each data point. Small values of *σ* produce “skinny” Gaussians that make every data point highly dissimilar from every other data point, and vice versa. Thus, *σ* effectively controls dimensionality of the Hilbert space induced by the kernel.

We implemented a heuristic reparameterization of MMD-MA that replaces the bandwidth parameter *σ* with a new parameter c. For a given collection of data points *X*^(1)^ and *X*^(2)^, we induce the corresponding latent representations *U*^(1)^ and *U*^(2)^ by using randomly initialized mapping matrices *α*_1_ and *α*_2_. We then compute Euclidean distances between all pairs *U_i_,U_j_* ∈ *U*^(1)^ ∪ *U*^(2)^, and we store the median Euclidean distance *d*. Intuitively, this value provides a numeric scale for the distances in the embedded space. We thus set the RBF kernel width as a constant factor of the scale d: *σ = cd*. We hypothesize that setting the bandwidth parameter in this data-driven way will allow us to fix *c*, thereby effectively reducing the number of hyperparameters from four to three. Note that *α*_1_ and *α*_2_ are initialized using values drawn from a uniform distribution over [0,1], and that these matrices also serve as the starting point for the MMD-MA optimization.

### 2.3 Datasets

Two sets of single-cell data sets from recent studies, each measuring both gene expression and chromatin accessibility, are used in our experiments.

#### SNARE-Seq data

Single-nucleus chromatin accessibility and mRNA expression sequencing (SNARE-seq) [11] is a recently described, droplet-based method that can link a cell’s transcriptome with its accessible chromatin for sequencing at scale. Because this method can measure both gene expression and chromatin accessibility information from the same cell, we use the SNARE-seq dataset to validate MMD-MA by evaluating the algorithm’s ability to correctly pair each expression profile with the correct chromatin accessibility profile. The cells assayed for this dataset consist of a mixture of BJ, H1, K562 and GM12878 cells, downloaded from the Gene Expression Omnibus database under accession number GSE126074.

Gene expression information in single cells is stored as cell × gene counts matrix *C_g_* of size *n*_1_ × *m*_1_, where *n*_1_ = 1047 is the number of cells with RNA-seq measurements and *m*_1_ = 18, 666 is the number of genes. The chromatin accessibility information is stored in a Boolean cell × peak matrix *C_a_* of size *n*_2_ × *m*_2_, where *n*_2_ = *n*_1_ = 1047 is the number of single cells with ATAC-seq measurements and *p*_2_ = 136, 771 is the number of peak regions. A value *C_a_*(*i,j*) = 1 indicates that peak *j* in cell *i* is accessible. As in the original SNARE-seq publication, we reduce data sparsity and noise in the ATAC-seq data by performing dimensionality reduction using the topic modeling framework cisTopic [12], resulting in a 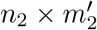 matrix with 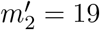

#### Mouse embryonic stem cell developmental data

The second dataset consists of sciRNA-seq [13] and sciATAC-seq [14] measurements, capturing transcription and chromatin accessibility information, respectively. These measurements were taken during embryoid body (EB) differentiation of mouse embryonic stem cells (mESCs) sampled at five stages: Days 0, 3, 7, 11, and finally as fully differentiated neural progenitor cells (NPCs).

This mESC line, female (F121-6), was derived from a cross between *Mus musculus* strain 129/SV-Jae (129) and *Mus castaneus* (cast) [15]. All mESCs were cultured and expanded on MEF feeders following the standard operating procedure designed for male mESC cell line F123 (4DN Consortium https://data.4dnucleome.org/protocols/1d39b581-9200-4494-8b24-3dc77d595bbb, except for the addition of 1% penicillin/streptomycin. Differentiation of mESCs into EBs was done using a standard LIF-withdrawal protocol. In brief, mESCs were grown on gelatin-coated plates for two passages to remove MEF feeders. Feeder-free mESCs were harvested and aliquots were saved as day 0 (D0). The remaining cells were cultured in regular medium (10% FBS and 1% penicillin/streptomycin in DMEM with high glucose) in non-adherent petri-dishes for 11 days of EB differentiation. EBs were collected at day 3, 7, and 11 (D3, D7 and D11) and treated with accutase to prepare single-cell suspensions for single-nucleus sciRNA-seq and sciATAC-seq. An aliquot of D11 EBs was used to derive NPCs using a published protocol [16]. NPCs were positively identified by staining with an antibody for nestin and used for sciRNA-seq and sciATAC-seq

Similar to the SNARE-seq data, the mouse expression and chromatin accessibility profiles are represented as two matrices *C_g_* and *C_a_*. However, for this dataset we reduced dimensions for both sciRNA-Seq and sciATAC-seq using cisTopic [12]. In this way, we obtained feature matrices *C_g_* of size *n*_1_ × *m*_1_, where *n*_1_ = 2127 and *m*_1_ = 60 and *C_a_* of size *n*_2_ × *m*_2_, where *n*_2_ = 1182 and *m*_2_ = 25.

Given that *n*1 ≠ *n*2, this dataset highlights the importance of developing unsupervised manifold alignment methods where it is hard or impossible to obtain one-to-one correspondence between the cells.

We perform *ℓ*_2_-normalization on the samples of the two datasets before converting them into scalar product similarity matrices *K*_1_ and *K*_2_.

### 2.4 Assessing alignment

For the SNARE-Seq dataset we have one-to-one correspondence information on the cell axis, because the data is captured from a single-cell co-assay. We do not provide this correspondence information to the MMD-MA algorithm, but instead use it to validate the performance of the method. To assess the alignment, we use the known correspondence between cells in the two domains as follows. For each cell x in one domain, we identify its true match in the other domain. We then rank all other-domain data points in the learned latent space by their distance from *x*, and we compute the fraction of points that are closer than the true match. Averaging this fraction across all data points in both domains yields the average “fraction of samples closer than the true match” (FOSCTTM), where perfect recovery of the true manifold structure will yield a value of zero.

For the mouse data, unlike the SNARE-Seq dataset, we do not have one-to-one correspondence information between the cells. However, we do have a time point label associated with each cell, corresponding to Days 0, 3, 7, 11, and NPCs. Therefore, to measure the performance of MMD-MA we evaluate how well the cells cluster in their respective time-point clusters after the alignment. We use the silhouette coefficient for this analysis [17], defined as (*b − a*)/ max(*a, b*), where *a* is the mean intra-cluster distance and *b* is the mean distance to nearest sample in a different cluster. The silhouette coefficient ranges from −1 (worst) to 1 (best). Since the silhouette coefficient represents the overall clustering quality, we join the two datasets and calculate the coefficient for the clusters of the combined data modalities.

### 2.5 Selection of hyperparameters

As described below, we fix one of MMD-MA’s four hyperparameters via simulations. Hence, our implementation of MMD-MA requires tuning the three remaining hyperparameters. To this end, we specified a hyperparameter grid as follows: weights λ_1_, λ_2_ ∈ {10^−3^,10^−4^,10^−5^,10^−6^,10^−7^} for the terms in the optimization problem and the dimensionality *p* ∈ {4, 5, 6} of the embedding space. To select hyperparameters we used the correspondence information and the time-point labels for the SNARE-seq and mouse datasets, respectively. We split the dataset into a 2:1 ratio, such that two-thirds of the samples were used as the training set and one-third as the validation set. Next, we ran the MMD-MA algorithm on the training dataset and calculated its performance on the validation set. We picked the set of hyperparameters that resulted in the lowest FOSCTTM score for SNARE-seq data and the highest silhouette score score for the mouse data. Finally, we ran the MMD-MA on the entire dataset using the selected hyperparameters and report the results in Section 3. This strategy allows us to choose the optimal hyperparameters in a systematic manner.

### 2.6 Baselines

We compare MMD-MA with the following unsupervised algorithms that do not require any correspondence information for aligning the datasets:

- **JLMA** [2] has one tunable hyperparameter, number of neighbors *k*, and we ran it for values *k* = 5 (default) and *k* = 6. For larger values of *k*, the algorithm runs too slowly to be practical.
- **UnionCom** [3] requires tuning of four hyperparameters and we specified the following search grid for them: the number *k* ∈ {5,10, 25, ...,*n*} (with increments of 25 after *k* = 25) of neighbors in the graph, the dimensionality *p* ∈ {3, 5,10} of the embedding space, the trade-off parameter *β* ∈ {0.001, 0.005, 0.01, 0.5, 0.1, 0.5,1, 5,10} for the embedding, and a regularization coefficient *ρ* ∈ {0.001, 0.005, 0.01, 0.5, 0.1, 0.5,1, 5,10}.
- **PCA** We used PCA to project the two datasets in the same space. This baseline demonstrates that there is little to no correspondence present in the data that can be easily extracted by performing dimensionality reduction and projection in the same space. We need to learn such correspondences by performing the alignment. Since PCA is indifferent to the orientation of each dimension, we treat the direction of each dimension as a hyperparameter. We present results for the optimally performing orientation.
- **Random Projection** We implemented another sanity check by randomly projecting the two datasets in the same space.

### 2.7 Implementation details

To allow the MMD-MA algorithm to scale up from the 61 cells used in the original publication to ~2000 analyzed here, we re-implemented the algorithm using Pytorch [18] to allow GPU usage. The Apache-licensed source code and documentation are available at https://bitbucket.org/noblelab/2020_mmdma_pytorch

## 3 Results

### 3.1 MMD-MA successfully integrates SNARE-seq data

To evaluate the performance of the MMD-MA algorithm, we begin by analyzing the SNARE-seq data. This data set contains gene expression and chromatin accessibility measurements which, when projected to 2D using PCA, exhibit qualitatively similar clusterings into three groups (Figure 2a–b). This is roughly similar to [11], where further pre-processing and t-SNE plots of the data result in four clusters representing BJ, GM12878, H1, and K562 cell mixtures. MMD-MA analysis of this data set, using a 5D latent space followed by projection to 2D via PCA, suggests that the algorithm succeeds in overlaying the two distributions (Figure 2c). This result is in contrast to UnionCom, which induces some shared structure but also groups a large cluster of gene expression measurements separately from any chromatin accessibility measurements (Figure 2d).

**Figure 2:**
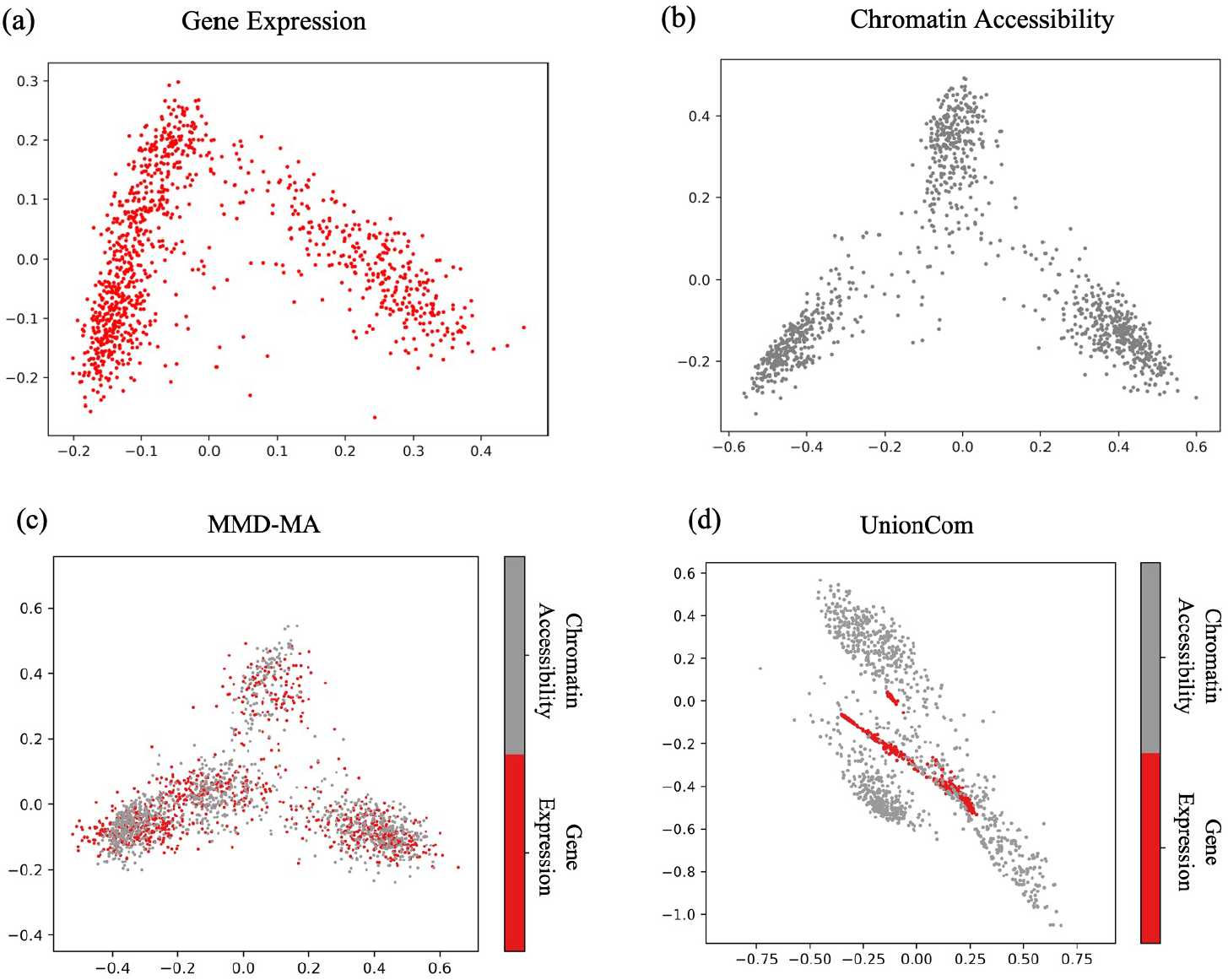
Integration of SNARE-seq data. PCA projection of the (a) gene expression and (b) chromatin accessibility profiles from the SNARE-seq data set. (c) The MMD-MA algorithm was used to project the SNARE-seq data into a 5-dimensional latent space, which was subsequently projected to 2D via PCA for visualization. (d) Projection for UnionCom (also reduced to 2D using PCA) for the best performing hyperparameters, using the reduced RNA-seq data (dimension *p* = 10).

To quantitatively evaluate these results, we exploit the fact that SNARE-seq is a co-assay and hence provides us with a true mapping between expression measurements and chromatin accessibility measurements. Using this mapping, we can compute the fraction of samples closer than the true match (FOSCTTM) score, as described in Methods. In addition to MMD-MA and UnionCom, we also apply JLMA and PCA to the data. All methods use a 5-dimensional latent space. We include as a negative control a random projection to 5 dimensions. The results (Figure 3) support the qualitative assessment from Figure 2, showing that MMD-MA achieves the lowest average FOSCTTM value of 0.16.

**Figure 3:**
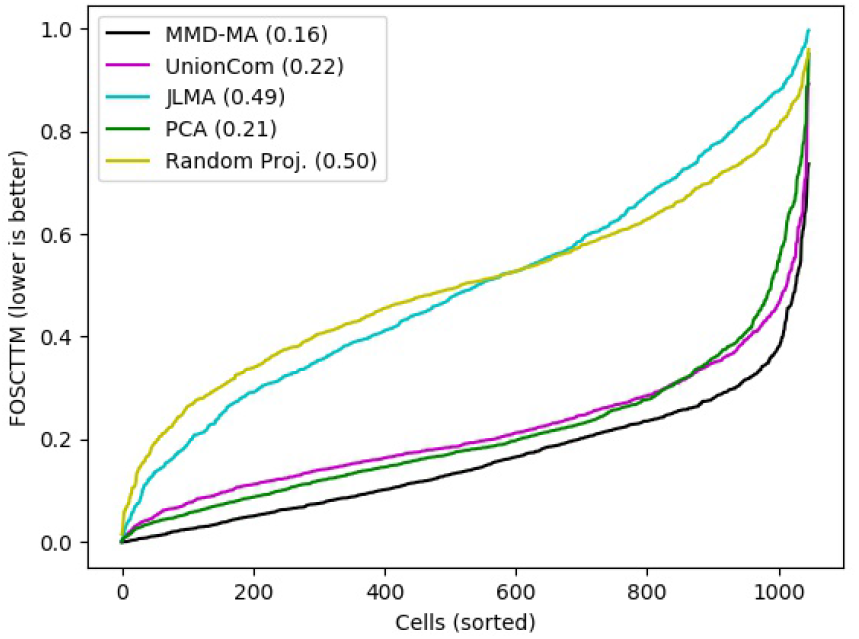
Comparison of MMD-MA’s performance on the SNARE-seq data with other unsupervised baselines. The figure plots the FOSCTTM score for all the cells in SNARE-Seq data, with cells sorted for each method by increasing score. Note that the UnionCom algorithm includes an initial preprocessing step, whereby the RNA-seq data is projected to 10 dimensions using PCA.

Note that UnionCom is too computationally expensive to run directly on the RNA-Seq measurements across 18,666 genes. Therefore, following [11], we first reduced the dimensions of the RNA-Seq measurements to 10 using PCA. To be sure that this PCA step is not the cause of the good relative performance of MMD-MA, we also ran MMD-MA on this 10-dimensional dataset. The performance (mean FOSCTTM of 0.14, Figure 4) is comparable to MMD-MA on the full dataset. Since UnionCom paper [3] does not specify data pre-processing normalization scheme, we applied both z-score and *l*_2_-normalization and found that the best performing score for z-score normalized data was higher (mean FOSCTTM of 0.26) than *l*_2_-normalized data (mean FOSCTTM of 0.22, presented in Figure 3). Both results were better than applying UnionCom on the datasets without normalization (mean FOSCTTM of 0.39) (See Figure 4).

**Figure 4:**
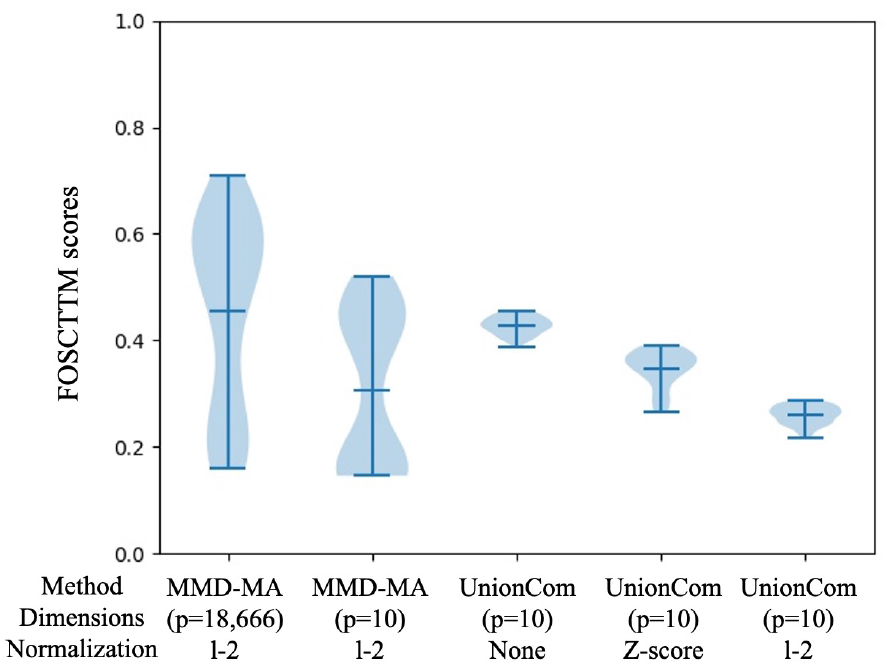
Comparison of MMDA-MA and UnionCom on the SNARE-seq data. The figure plots the distribution of FOSCTTM scores across a grid of hyperparameters for MMD-MA and UnionCom. Two variants of MMD-MA are included, one that operates on the original RNA-Seq count matrix with feature dimension *p* = 18,666, and one that operates on RNA-seq data that has been reduced to *p* = 10 dimensions using PCA. Since UnionCom paper does not specify data pre-processing normalization scheme, we applied both z-score and *l*_2_-normalization and found that the best performing score for z-score normalized data was higher (mean FOSCTTM of 0.26) than *l*_2_-normalized data (mean FOSCTTM of 0.22). Both results were better than applying UnionCom on the datasets without normalization (mean FOSCTTM of 0.39). Overall, the best performing FOSCTTM scores obtained for MMD-MA (0.16 and 0.14) are much lower than that of UnionCom.

### 3.2 Accurate time-point based clustering of mouse embryonic stem cell data

We next turn to the mouse embryonic stem cell time series data. Both components of this dataset, gene expression and chromatin accessibility, exhibit a boomerang-shaped structure when visualized using PCA, corresponding to the developmental trajectory (Figure 5a–b). Among the five time points, the neural precursor cells (NPCs) appear to be the most distinct in both data modalities.

**Figure 5:**
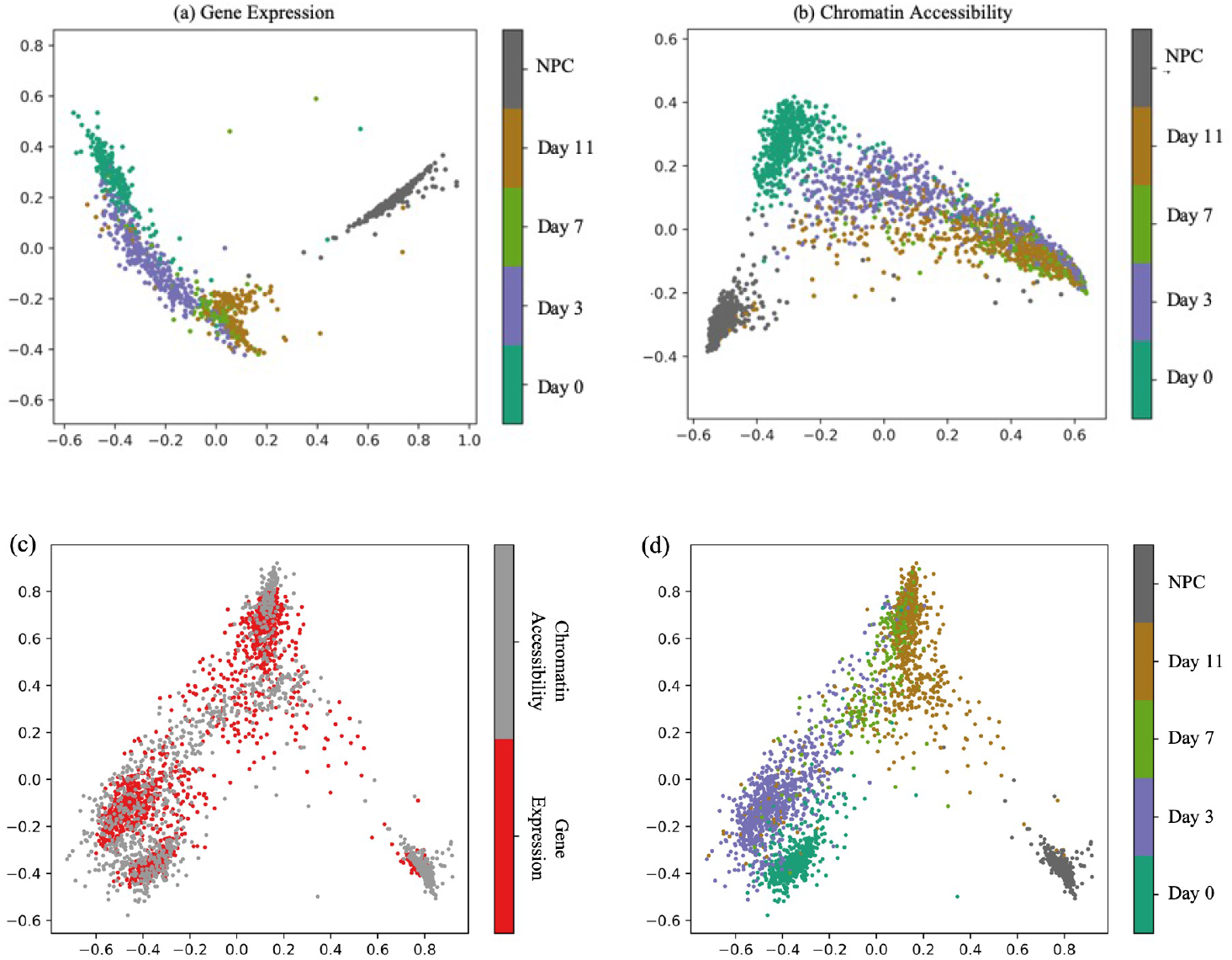
Mouse embryonic stem cell data. (a) The first two principal components of the mouse scRNA-Seq data. (b) Similar to panel (a), but for the scATAC-seq data. (c) The 4-dimensional embedding learned by MMD-MA, projected to 2D via PCA. Points are colored according to data type. (d) Same as panel (c), but coloring points by time point.

Projection of the data to five dimensions, followed by 2D PCA visualization, suggests that MMD-MA successfully overlays the two data sets (Figure 5c), with measurements from the same developmental time point lying near one another in the latent space (Figure 5d). To quantify this agreement, we compute the silhouette coefficient for each time point. The resulting values agree with the visual layout in Figure 5d: the NPC time point gives the best alignment performance of 009323, whereas the Day 7 and 11 clusters, which overlap in both the expression and chromatin accessibility projections, achieve the lowest silhouette coefficients (0.00214 and −0.00525, respectively).

We also use the silhouette analysis to quantitatively compare MMD-MA to UnionCom, JLMA, PCA and random projections (Table 2). Note that this comparison is slightly unfair, in favor of UnionCom: for UnionCom we report the results from the best performing hyperparameter values, whereas for MMD-MA we pick hypeparameters using cross-validation, as described in Methods. Nonetheless, MMD-MA’s average silhouette coefficient is the highest among all methods. Furthermore, the silhouette coefficient for MMD-MA alignment is consistently better than JLMA for all five timepoints and better than UnionCom in four out of five cases. In general, all methods struggle to discriminate between data from days 7 and 11, which appear to be quite similar to one another.

**Table 2:** Comparison of MMD-MA with other unsupervised baselines using the mouse data set. The table lists the silhouette score for each method with respect to each time point in the mouse embryonic stem cell data set, with the average score in the final column. The best-performing method in each column is indicated in boldface.

| Method | Day 0 | Day 3 | Day 7 | Day 11 | NPC | Avg. |
| --- | --- | --- | --- | --- | --- | --- |
| MMD-MA | **0.7633** | **0.5107** | 0.0214 | **-0.0525** | **0.9323** | **0.4767** |
| UnionCom | 0.6906 | 0.4788 | **0.0824** | -0.1025 | 0.8076 | 0.4223 |
| JLMA | 0.1674 | -0.0691 | -0.3229 | -0.3684 | 0.6019 | 0.0239 |
| PCA | 0.4355 | 0.4661 | -0.055 | -0.1239 | 0.6059 | 0.3095 |
| Random Projection (avg.) | 0.2825 | -0.0031 | -0.1762 | -0.4136 | 0.6997 | 0.0887 |

To investigate whether integration of the two data modalities leads to improved clustering within each data set, we compute silhouette coefficients separately within each data modality (sciRNA-seq and sciATAC-seq, Figure 6). Not surprisingly, projecting to 5D using PCA consistently improves the silhouetted coefficient relative to analysis carried out in the original data space. This observation holds across all time points and across both data modalities. However, when we compare PCA, which is performed independently on the two data modalities, to MMD-MA, which is performed jointly, we see a consistent improvement in the silhouette scores computed with respect to the sciATAC-seq data. We hypothesize that during the alignment step, information from the sciRNA-seq data perturbs the sciATAC-seq measurements to better local clustering. The converse is not true—sciRNA-seq clustering is approximately the same using PCA or MMD-MA—presumably because the sciRNA-seq data is less noisy, more informative with respect to developmental time, or both.

**Figure 6:**
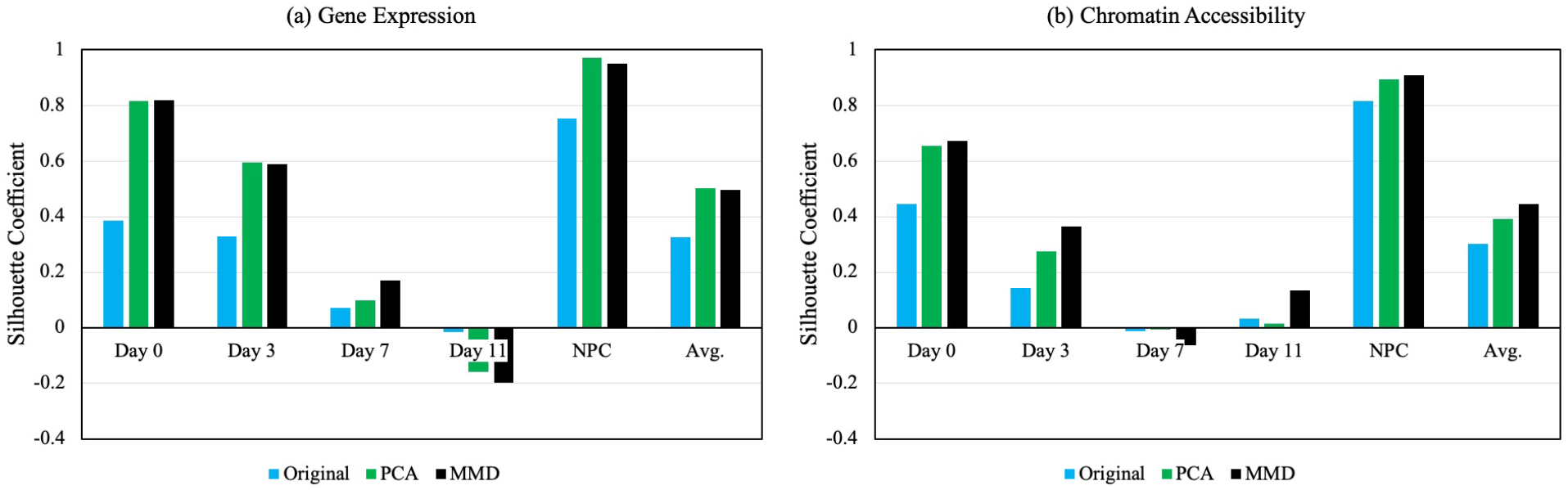
Evaluation of the time-point based clustering within each sciRNA-seq and sciATAC-seq dataset. MMD-MA not only preserves the local structure of the two datasets, but improves the clustering performance of sciATAC-Seq data. We hypothesize that during the alignment step, the time-point information from sciRNA-seq guides the cell measurements of sciATAC-seq to better local clustering.

### 3.3 Improved implementation

#### Faster GPU computation

To compare the running time of our MMD-MA GPU implementation with the original one in [9], we subsampled 50, 100, 200, 400, and 800 cells from the SNARE-Seq dataset and ran both implementations of MMD-MA. We ran the GPU implementation on an Nvidia GTX 1080, and the CPU implementation on an AMD Athlon^*™*^ × 4950. Our GPU implementation scales well with the number of cells (Figure 7a), requiring ~ 4 minutes to run 20,000 iterations of MMD-MA on 1047 cells, whereas the CPU implementation takes ~ 100 minutes for the same task.

**Figure 7:**
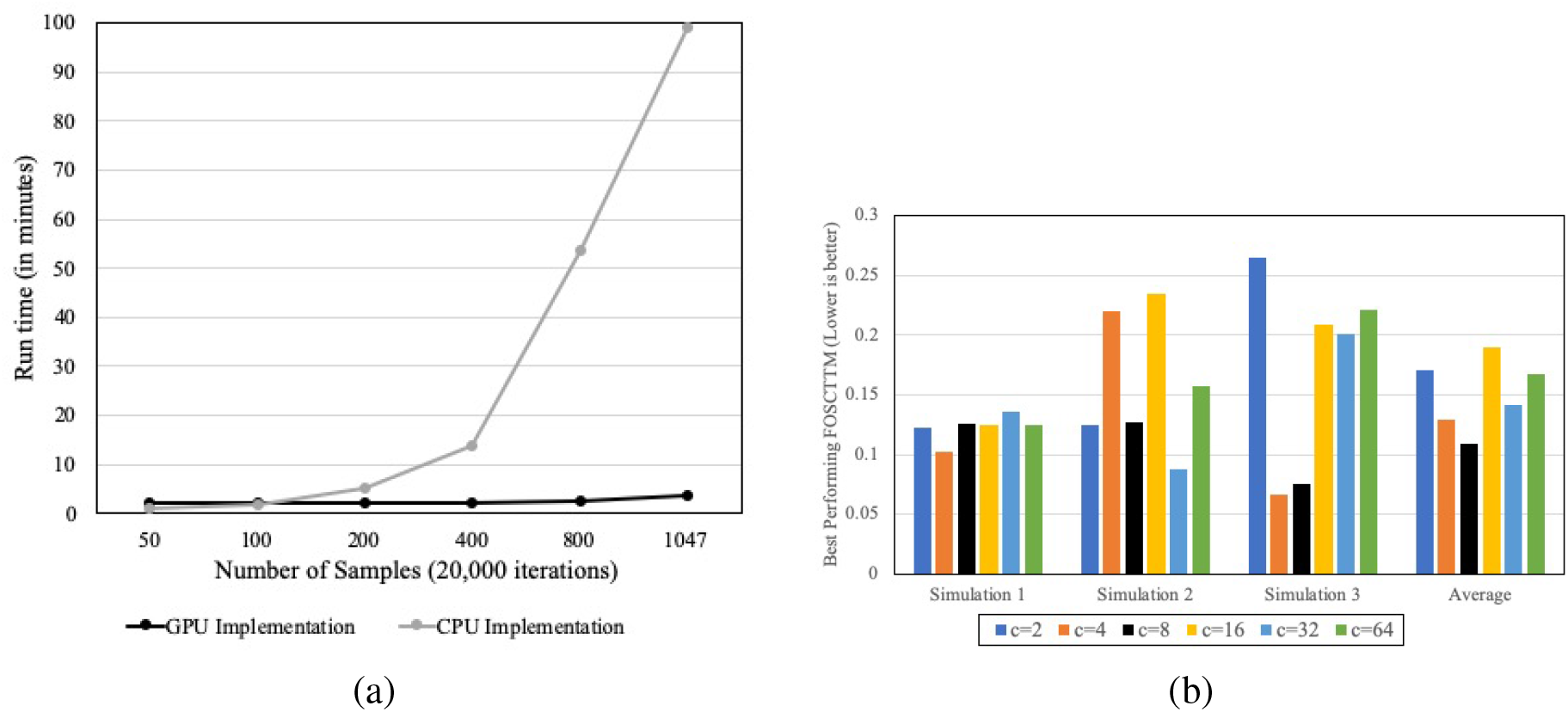
Improved MMD-MA implementation. (a) The figure plots wall clock time as a function of the number of cells, with series corresponding to the original CPU implementation of MMD-MA and our GPU implementation. (b) Best performing FOSCTTM values for running the reparameterized MMD-MA on the three recreated simulations from [9] for different values of *c*.

#### Automatic bandwidth selection

We reparameterize MMD-MA by replacing the bandwidth parameter *σ* with a new parameter c. We aim to fix *c* and reduce the effective number of hyperparameters from four to three. To select a value of *c*, we re-created the three simulation datasets from [9] with a larger number of samples (*n* = 1000). The three simulations consist of points belonging to a shared manifold (branch, swiss-roll, and frustum) and projected non-linearly to a high dimensional space. The number of samples was increased to represent the sample size of our real-world datasets. Given that all the points have correspondence information, we can use the FOSCTTM value to evaluate the alignment performance. Therefore, we ran our reparameterized MMD-MA algorithm that calculated *σ* with different values of *c* ∈ {2, 4, 8,16, 32, 64} and recorded the best performing FOSCTTM values for each.

As shown in Figure 7b, we observe that different values of *c* give widely varying performance. For example, FOSCTTM values for *c* = 4 range from 0.10 (simulation 1) to 0.22 (simulation 2) to 0.06 (simulation 3). However, *c* = 8 consistently gives low FOSCTTM values across all three simulations with minimal variation (standard deviation OoO29). This is summarized by taking the averages of the FOSCTTM values across the three simulations. Based on these results, we fixed the parameter as *c* = 8 in the analyses of SNARE-seq and mouse data reported above.

## 4 Discussion

Integration of heterogeneous single-cell data sets requires the development of effective alignment algorithms. We demonstrate that MMD-MA can be effectively used to align real-world single-cell experiments by applying it to two recent state-of-the-art datasets that measure single-cell transcriptome and chromatin accessibility. We evaluate MMD-MA’s alignment performance and show that it outperforms existing state-of-the-art unsupervised alignment methods.

We extend the MMD-MA implementation to GPUs, allowing it to scale to thousands of samples and improve its running speed by a factor of 25. Finally, we also reduce the hyperparameter search space for the MMD-MA algorithm by proposing a method to automatically select the *σ* parameter that determines the bandwidth during kernel calculation.

We acknowledge that we will not be able to use hyperparameter tuning for datasets without available labels or correspondence information (that is, in a truly unsupervised setting). Therefore, analyzing hyperparameters and devising strategies to select them automatically is a part of the future work for this study.

Our approach of automatically assigning the *σ* parameter (Section 2.2) is the first step in that direction. For now, a reduced hyperparameter search space and a fast GPU-based framework allow our MMD-MA implementation to serve as a useful tool to perform single-cell alignment of different real-world datasets.

## References

[1] T. Stuart and R. Satija. Integrative single-cell analysis. Nature Reviews Genetics, 20:252–272, 2019.

[2] C. Wang, P. Krafft, and S. Mahadevan. Manifold alignment. In Y. Ma and Y. Fu, editors, Manifold Learning: Theory and Applications. CRC Press, Boca Raton, FL, USA, 2011.

[3] K. Cao, X. Bai, Y. Hong, and L. Wan. Unsupervised topological alignment for single-cell multi-omics integration. bioRxiv, 2020. https://doi.org/10.1101/2020.02.02.931394.

[4] Z. Cui, H. Chang, S. Shan, and X. Chen. Generalized unsupervised manifold alignment. In Z. Ghahramani, M. Welling, C. Cortes, N. D. Lawrence, and K. Q. Weinberger, editors, Advances in Neural Information Processing Systems 27, pages 2429–2437. Curran Associates, Inc., Montreal, Canada, 2014.

[5] J. D. Welch, A. J. Hartemink, and J. F. Prins. MATCHER: manifold alignment reveals correspondence between single cell transcriptome and epigenome dynamics. Genome biology, 18(1):138, 2017.

[6] M. Amodio and S. Krishnaswamy. MAGAN: Aligning biological manifolds. In Jennifer Dy and Andreas Krause, editors, Proceedings of the 35th International Conference on Machine Learning, volume 80 of Proceedings of Machine Learning Research, pages 215–223, Stockholmsmässan, Stockholm Sweden, 10-15 Jul 2018. PMLR.

[7] J. D. Welch, V. Kozareva, A. Ferreira, C. Vanderburg, C. Martin, and E. Z. Macosko. Single-cell multi-omic integration compares and contrasts features of brain cell identity. Cell, 177(7):1873–1887, 2019.

[8] T. Stuart, A. Butler, P. Hoffman, C. Hafemeister, E. Papelexi, W. M. Mauck III, Y. Hao, M. Stoeckius, P. Smibert, and R. Satija. Comprehensive integration of single-cell data. Cell, 77(7):1888–1902, 2019.

[9] J. Liu, Y. Huang, R. Singh, J.-P. Vert, and W. S. Noble. Jointly embedding multiple single-cell omics measurements. In Katharina T. Huber and Dan Gusfield, editors, 19th International Workshop on Algorithms in Bioinformatics (WABI 2019), volume 143 of Leibniz International Proceedings in Informatics (LIPIcs), pages 10:1–10:13, Dagstuhl, Germany, 2019. Schloss Dagstuhl-Leibniz-Zentrum fuer Informatik.

[10] N. Cristianini and J. Shawe-Taylor. An Introduction to Support Vector Machines. Cambridge UP, Cambridge, UK, 2000.

[11] S. Chen, B. B. Lake, and K. Zhang. High-throughput sequencing of the transcriptome and chromatin accessibility in the same cell. Nature Biotechnology, 37(12):1452–1457, 2019.

[12] C. B. González-Blas, L. Minnoye, D. Papasokrati, S. Aibar, G. Hulselmans, V. Christiaens, K. Davie, J. Wouters, and S. Aerts. cisTopic: cis-regulatory topic modelling on single-cell ATAC-seq data. Nature Methods, 16(5):397–400, 2018.

[13] J. Cao, J. S. Packer, V. Ramani, D. A. Cusanovich, C. Huynh, R. Daza, X. Qiu, C. Lee, S. N. Furlan, F. J. Steemers, et al. Comprehensive single-cell transcriptional profiling of a multicellular organism. Science, 357(6352):661–667, 2017.

[14] D. A. Cusanovich, J. p. Reddington, D. A. Garfield, R. M. Daza, D. Aghamirzaie, R. Marco-Ferreres, H. A. Pliner, L. Christiansen, X. Qiu, F. J. Steemers, et al. The cis-regulatory dynamics of embryonic development at single-cell resolution. Nature, 555(7697):538, 2018.

[15] H. Marks, H. H. D. Kerstens, T. S. Barakat, E. Splinter, R. A. M. Dirks, G. van Mierlo, O. Joshi, S. Wang, T. Babak, C. A. Albers, T. Kalkan, A. Smith, A. Jouneau, W. de Laat, J. Gribnau, and H. G. Stunnenberg. Dynamics of gene silencing during x inactivation using allele-specific rna-seq. Genome Biology, 16(1):149, 2015.

[16] H. Xu, X. Fan, J. Tang, G. Zhou, L. Yang, X. Wu, S. Liu, J. Qu, and H. Yang. A modified method for generation of neural precursor cells from cultured mouse embryonic stem cells. Brain research protocols, 15(1):52–58, 2005.

[17] P.J. Rousseeuw. Silhouettes: a graphical aid to the interpretation and validation of cluster analysis. Journal of Computational and Applied Mathematics, 20:53–65, 1987.

[18] A. Paszke, S. Gross, F. Massa, A. Lerer, J. Bradbury, G. Chanan, T. Killeen, Z. Lin, N. Gimelshein, L. Antiga, A. Desmaison, A. Kopf, E. Yang, Z. DeVito, M. Raison, A. Tejani, S. Chilamkurthy, B. Steiner, L. Fang, J. Bai, and S. Chintala. Pytorch: An imperative style, high-performance deep learning library. In Advances in Neural Information Processing Systems 32, pages 8024–8035. Curran Associates, Inc., Vancouver, Canada, 2019.

